# The representation of stimulus features during stable fixation and active vision

**DOI:** 10.1101/2024.10.09.617414

**Authors:** Caoimhe Moran, Philippa A. Johnson, Hinze Hogendoorn, Ayelet N. Landau

## Abstract

Predictive updating of an object’s spatial coordinates from its retinotopic pre-saccadic to post-saccadic position contributes to stable visual perception. However, whether object features are predictively represented at the remapped location remains contested. Many previous studies showing evidence of feature remapping neglect the spatially invariant representation of features in the visual system. For example, feature-based attention boosts attended features across the entire visual field, potentially contributing to the maintenance of stimulus features across saccades. We set out to characterise the spatiotemporal dynamics of feature processing during stable fixation and active vision. To do so, we applied multivariate decoding methods to electroencephalography (EEG) data collected while participants viewed brief visual stimuli. Stimuli appeared at different locations across the visual field at either high or low spatial frequency (SF). During fixation, classifiers were trained to decode SF presented at one parafoveal location and cross-tested on SF from either the same, adjacent or more peripheral locations. When training and testing on the same location, SF was classified shortly after stimulus onset (∼80 ms). Decoding of SF at locations farther from the trained location emerged later (∼150-300 ms), with decoding latency modulated by eccentricity. This analysis provides a detailed time course for the spread of feature information across the visual field. Next, we investigated how active vision impacts the emergence of SF information. In the presence of a saccade, the decoding time of peripheral SF at parafoveal locations was accelerated, indicating predictive anticipation of SF due to the saccade. Crucially however, this predictive effect was not limited to the specific remapped location. Rather, peripheral SF was correctly classified, at an accelerated time course, at all parafoveal positions. This indicates a spatially coarse remapping of stimulus features during active vision, likely enabling a smooth transition on saccade landing.

**Significance Statement:** Maintaining a continuous representation of object features across saccades is vital for stable vision. In order to characterise the spatiotemporal dynamics of stimulus feature representation in the brain, we presented stimuli at a high and low spatial frequency at multiple locations across the visual field. Applying EEG-decoding methods we tracked the neural representation of spatial frequency during both stable fixation and active vision. Using this approach, we provide a detailed time course for the spread of feature information across the visual field during fixation. In addition, when a saccade is imminent, we discovered that the spread of feature information is expedited such that access to peripheral spatial frequency is accelerated.

## 1 Introduction

The early visual system encodes the world retinotopically: objects are projected to early visual areas with spatial coordinates determined by their position on the retina. This complicates accurate localisation, given that eye movements continuously shift the retinal position of objects. Despite this, visual perception remains largely unperturbed. There is a general consensus that predictive remapping of a stimulus’ spatial coordinates from their retinotopic pre-saccadic to post-saccadic position contributes to visual stability (Mays & Sparks, 1980; Bruce & Goldberg, 1985; Duhamel et al., 1992; Walker et al., 1995; Thompson et al., 1996; Umeno & Goldberg, 1997; Gottlieb et al., 1998; Nakamura & Colby, 2002; Sommer & Wurtz, 2002; Merriam et al., 2007; Parks & Corballis, 2008; Knapen et al., 2016; Golomb, 2019; Moran et al. 2024).

Spatial remapping has been demonstrated in animal neurophysiology (Bruce & Goldberg, 1985; Duhamel et al., 1992; Gottlieb et al., 1998; Kusunoki et al., 2000; Nakamura & Colby, 2002; Umeno & Goldberg, 2001), human behaviour (Hunt & Cavanagh, 2011) and human neuroimaging (Merriam et al., 2003, 2007; Medendorp et al., 2005, Parks & Corballis, 2010, Moran et al. 2024). Neurons fire predictively in response to a stimulus presented outside their classical receptive field, in a location corresponding to the future field of that neuron (i.e., the area of space encompassed by the receptive field post-saccade). We recently investigated the temporal unfolding of spatial remapping in humans (Moran et al. 2024). Using EEG in combination with multivariate pattern analysis to provide high temporal precision, we found that evidence for the stimulus at the remapped location could only be found when the stimulus was presented in a short time window immediately before saccade onset. Stimuli presented with longer latencies before the saccade were processed at their veridical retinotopic location. In the brief window immediately before a saccade, the brain is in a state of motor preparation, implying that the corollary discharge signal plays a role in activating receptive fields overlapping the remapped location. Our results reveal the time course of spatial remapping, finding above-chance decoding of the remapped position at 180 ms post-stimulus onset, and highlight the importance of the relative timing between stimulus and saccade.

However, visual stimuli are defined by more than their spatial location. It is well documented that the visual system also processes low-level (e.g., orientation, spatial frequency, colour) (Enroth-Cugell et al., 1983; Orban et al., 1984; Ringach et al., 2003; Watson et al., 2016) and high-level stimulus features (e.g. faces, categories) (Kiani et al., 2005; Rossion & Jacques, 2011) in dedicated neuronal populations throughout the visual hierarchy (Hubel & Wiesel, 1969; Zeki, 1978; Albright, 1984). If the role of remapping is to ensure the continuity of visual input across a saccade, then anticipation of the relevant spatial coordinates is not enough. With each saccade, the projection of visual stimuli on the retina undergoes a major transformation, such that, on saccade landing, stimuli are processed by different sets of neurons than before the saccade. If only spatial information is remapped then post-saccadic neurons would be required to re-process stimulus features from scratch with each new fixation, as they would have no information about its identity. For accurate visual perception across saccades, it is important that we can track input and also identify changes in that input. To achieve this, some information about stimulus identity could be relayed from pre-saccadic to post-saccadic neurons (Hollingworth et al., 2008; Gordon et al., 2008; Melcher & Colby, 2008). Whether saccadic remapping is a feature selective process is a highly contentious issue.

Some researchers argue in favour of a transfer of features including orientation (He et al., 2018) and letters (Harrison et al., 2013) around the time of a saccade. In contrast, others have suggested that remapping is a purely spatial process and additional mechanisms such as modulatory feedback are recruited to maintain stimulus identity and link it to spatial location after the saccade (Cavanagh et al., 2010). The majority of studies investigating the transfer of featural information across saccades in humans are behavioural. For example, Harrison et al., (2013) exploited characteristics of visual crowding to examine whether distractors presented at a stimulus’ post-saccadic retinotopic position would interfere with identification of a letter presented in the periphery. They showed that, in contrast to a fixation condition, identification of the letter was significantly impaired around the time of a saccade when the stimulus was remapped to a position surrounded by distractors. Other influential studies utilised the tilt after-effect (TAE; Gibson & Radner, 1937; Gibson, 1937) to show that illusory effects were preserved across saccades (Melcher,2005, 2009; Zimmermann et al., 2017) suggesting a transfer of features to the post-saccadic retinotopic location when a saccade is imminent. A major issue with these studies however is the limited sampling of spatial locations, making it difficult to determine if TAE and crowding effects are unique to the remapped location or more widespread. Studies investigating the motion after-effect and TAE outside of the saccade literature have found that an adaptor presented at one location induced an after-effect evenly across an array of locations tested (Liu & Hou, 2011; Liu & Mance, 2011). This was in the absence of any saccade and so may reflect properties of feature-based attention (FBA), which acts across the entire visual field.

To rule out FBA as an explanation for the influence of peripheral stimuli on perception at the remapped location, a recent study investigating feature remapping included a no-saccade condition (Fabius et al., 2019). If perceptual alterations at the remapped location are due to stimulus remapping then no effect should be found in the absence of a saccade. Employing the hi-phi illusion (Wexler et al., 2013), researchers found that, during stable fixation a peripheral inducer lead to illusory effects at fixation (the remapped location). However, the effect was significantly weaker compared to when the inducer was viewed during saccade preparation. This indicates that while spatially invariant processes may explain a small proportion of the effect, there is evidence that as a result of the upcoming saccade, the pre-saccadic inducer is remapped and thus influences perception of the stimulus on saccade landing, explaining the enhanced illusory effect. However, a major limitation in this study and others (Herwig & Schneider, 2014; Herwig, 2015; Paeye et al., 2017) is that the stimulus to be remapped is the saccade target, meaning the remapped location overlaps with the fovea. This makes it difficult to determine if the effect can be attributed to feature remapping or a fovea-specific effect, such as foveal feedback. There is plenty of evidence from psychophysics (Fan et al, 2016; Yu & Shim, 2016 & Weldon et al., 2020), neuroimaging (Williams et al. 2008; Fan et al., 2016) and brain stimulation (Chambers et al., 2013) to suggest that the fovea contributes to the processing of peripheral stimuli during stable fixation. For example, using functional magnetic resonance imaging, Williams et al., (2008) found that the pattern of fMRI activity at foveal retinotopic cortex contained category information about peripheral stimuli in the absence of a saccade. Recent behavioural research into foveal feedback under saccade conditions found that participants had increased sensitivity to orientations presented in foveal regions when they matched the saccade target (Kroell & Rolfs, 2022). Remapping was ruled out as an explanation for the results as enhancement of saccade target features better aligned to the centre of gaze than the remapped target position, determined based on the saccade endpoint (Collins et al., 2009).

Invasive neurophysiological recordings in animals and neuroimaging methods in humans have demonstrated that neurons in LIP, SC, FEF and extra striate cortex exhibit remapping characteristics (Duhamel et al., 1992; Walker et al., 1995; Umeno & Goldberg, 1997; Nakamura & Colby, 2002; Merriam et al., 2003, 2007; Zirnsak & Moore, 2014; Wang et al., 2023). Despite this, there is almost no direct neural evidence that the responses recorded at remapped locations contain information about stimulus features. There are two exceptions that offer conflicting results. One animal study demonstrated shape selectivity in the future field of neurons in LIP (Subramanian & Colby, 2014) while another study, recording from area MT in monkeys, found no selectivity for motion direction in the remapped response (Yao et al., 2016). There is even less neurophysiological evidence in humans. That said, a recent study investigated whether spatial frequency (SF) was updated across a saccade (Fabius et al., 2020). They did not find pre-saccadic remapping of SF and instead report that both the pre-saccadic and post-saccadic representation of SF could be decoded from MEG after saccade landing. This means that for approximately 200 ms after saccade landing, both stimulus representations are accessible, before the post-saccadic stimulus prevails. As in previous studies, only one location was sampled and the stimulus remained on the screen throughout the entire trial, limiting the conclusions we can draw about a predictive representation of SF.

Given previous evidence that peripheral stimulus information can influence perception and the neural response at unstimulated positions, even in the absence of a saccade, it is important to understand the time course of this process. This can be done by taking a temporally resolved approach to the spread of stimulus information across the visual field. This allows a dissociation between information at the remapped location due to feedback processes that exist even in the absence of a saccade and information transferred to the remapped location as a result of saccadic remapping. In addition, to circumvent confounding fovea-specific effects with effects unique to remapping, the stimulus to be remapped needs to be separated from the saccade target, inducing peripheral to peripheral remapping rather than peripheral to foveal remapping. Finally, to ensure bottom-up visual input does not interfere with the unfolding dynamics of the remapped representation, a brief stimulus must be used so that no visual input enters the eye at the post-saccadic retinotopic position.

To investigate the remapping of stimulus features across a saccade and determine the spatial specificity of the remapped response, we conducted a decoding analysis of human EEG data sampling multiple stimulus locations. Participants completed fixation trials and saccade trials during which a grating stimulus, at a high or low SF, was briefly presented at various locations on the screen. Before investigating whether SF shifted to the remapped location as a result of the saccade, we were interested in the temporal unfolding of the representation of SF during stable fixation. To this end, we sought to decode SF information from each stimulus location while participants maintained fixation. To characterise the extent to which SF information spreads across the visual field, we ran a cross-location decoding analysis, training a classifier on a single location and testing it on other locations. To foreshadow the results, we found that SF information spreads from the actual stimulus position across the cortex as a function of eccentricity, ultimately providing a position invariant representation of SF. Secondly, we aimed to determine whether SF is predictively remapped to the post-saccadic retinotopic position. For this, we again performed a cross-location decoding analysis, training on fixation trials and testing on saccade trials. We found that the SF of peripheral stimuli in saccade trials could be decoded earlier at the post-saccadic retinotopic location than stimuli at the same eccentricity during stable fixation. Importantly however, by testing incongruent locations we found that this early decoding is not unique to the remapped location but rather signifies a spatially coarse remapping of SF to positions close to fixation.

## 2 Methods

Participants, experimental design and data pre-processing were adapted from Moran, Johnson, Landau and Hogendoorn, (2024), and existing data from this paper was re-analysed here.

### 2.1 Participants

Participants were required to complete six testing sessions across different days, including one screening session. Thirteen observers took part in the initial screening session, of whom three were excluded due to poor EEG classification performance (less than 52% average decoding accuracy when classifying stimulus presentation location on fixation trials). After exclusion, ten observers (three male; mean age = 25.7 years, sd = 2.0 years) with normal or corrected-to-normal vision remained. The experimental protocol was approved by the human research ethics committee of The University of Melbourne, Australia (Ethics ID: 2021-12985-16726-4) and conducted in accordance with the Declaration of Helsinki. All observers provided informed consent before beginning the experiment and were reimbursed AU$15 per hour for their time, plus an additional AU$20 after completing all sessions.

### 2.2 Stimuli and procedure

Stimuli were programmed in MATLAB Version R2020a, using the Psychophysics Toolbox extension (Brainard, 1997; Pelli, 1997; Kleiner, Brainard & Pelli, 2007). They were presented on an ASUS ROG PG258 monitor (ASUS, Tapei, Taiwan) with a resolution of 1,920 x 1,080 running at a refresh rate of 120 Hz. Participants were seated in a quiet, dark room with their head supported by a chin rest, positioned 80 cm from the screen.

Stimuli consisted of sinusoidal gratings presented within a circular Gaussian window (i.e. Gabors; outer diameter: 7.9 degrees of visual angle (dva); 100% contrast) presented on a grey background for 100 ms. The stimulus could appear at a SF of .33 c/dva or 1 c/dva while the orientation of the grating was fixed at 0⁰ (except on catch trials). For catch trials, participants were instructed to respond to an oddball grating of a different orientation (90⁰) which could appear every 11-20 trials. These trials were included in order to ensure attentive processing of the grating stimuli, and were excluded from the main analysis.

Two fixation points (.41 dva), one black and one white, were horizontally aligned and subtended 10.04 dva to the left and right of the screen centre (20.08 dva apart). They appeared at the beginning of the experiment and remained visible throughout, excluding breaks. Participants were instructed to fixate on the black fixation point at all times.

Periodically during the experiment, a saccade cue appeared: the colour of the fixation points gradually changed, such that over a period of 1.2s the black fixation point became white and vice versa. Participants were instructed to monitor the colour of the fixated fixation point, and plan and execute a saccade from one fixation point to the other as soon as they detected the colour change, such that they were always fixating the black fixation point.

Different conditions were determined by the conjunction of fixation position and the location of the grating stimulus. In fixation trials, the fixation was stable (no colour change occurs), and stimuli appeared in one of four positions around the current black fixation point (location 1-4; Fig. 1a. *right*). In control trials the fixation was stable, and stimuli appeared on the other side of the screen, in one of four locations around the white fixation point (location 5-8; Fig. 1a. *right*). In saccade trials, there was a colour change and stimuli appeared during the planning and execution of a saccade. In this case, stimuli could appear around current fixation or saccade target (saccade trials) (location 5-8; Fig. 1a *right*) (see Fig. 1 b and c for task design).

**Figure 1:**
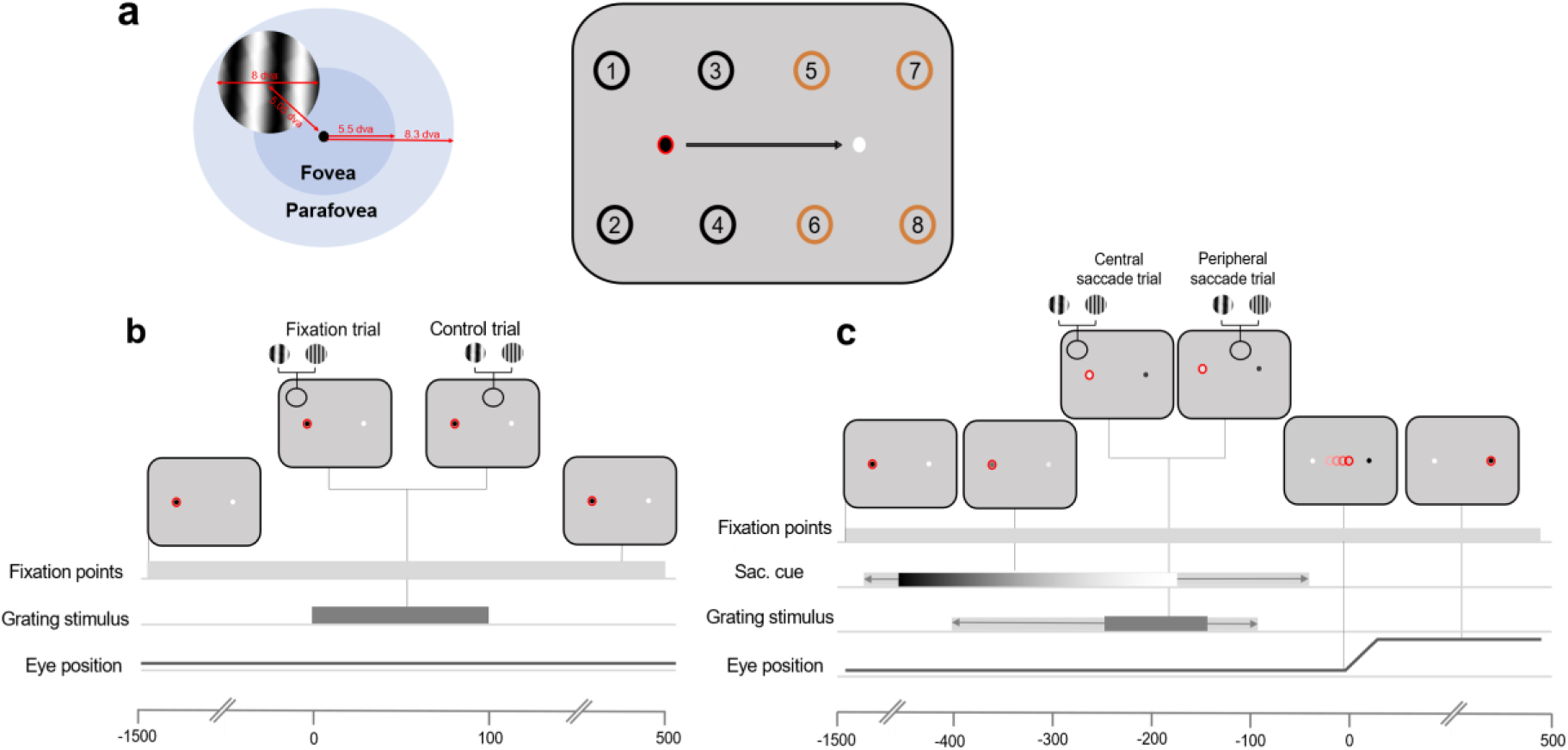
adapted from Moran et al., 2024. a. Stimulus dimensions and configuration. *Left*: Stimuli were sinusoidal gratings (diameter, 7.9 dva) presented 5.05 dva from a fixation point. Stimuli extended across foveal and parafoveal regions. *Right:* Participants always fixated on the black fixation point and made a saccade when cued by a gradual colour change to white. Stimuli were presented around fixation on fixation trials (location 1-4; black) or around the white fixation point/saccade target on control trials and saccade trials, respectively (location 5-8; orange). The red circle around the black fixation point indicates eye position. The numbers on the screen indicate all possible stimulus presentation locations. **b.** In fixation trials, participants maintained fixation on the black fixation target (shown here with a red ring to indicate the position of the eyes) throughout the trial, never fixating the white. Approximately 400-800 ms after trial onset, the grating stimulus was presented at one of four locations around fixation at either a low or high SF. The stimulus was displayed for 100 ms. In control trials, participants fixated on the black fixation point. A grating stimulus was presented at one of four locations around the alternate fixation point (white circle). **c.** In saccade trials, participants fixated on the black fixation point. After 200-600 ms the black fixation point changed to white and the white fixation point changed to black, which served as the saccade cue. The colours gradually changed over a period of 1.2 s, indicated here by the colour gradient. The light grey bar behind the gradient refers to possible start/end times of the colour change. In a period ranging from 400 ms to 100 ms before saccade onset, a grated stimulus was presented at one of four locations around the saccade target. The stimulus was presented for 100 ms. The light grey bar behind dark grey bar refers to possible start/end times of the stimulus.

Stimulus locations were separated by 5.06 dva and were also 5.06 dva away from the nearest fixation point. The fovea covers the central 5.5 dva while the parafovea covers 8.3 dva around fixation (Hendrickson, 2005). Our grating stimuli covered 8 dva and therefore, on fixation trials, extended across foveal and parafoveal regions (Fig. 1a. *left*). For simplicity, we refer to these stimuli as parafoveal stimuli. We discuss the implications of this stimulus placement in the discussion section.

The grating was presented briefly at a variable delay after the saccade cue. This delay was adjusted for each participant in order to sample trials with approximately 200 ms between stimulus onset and saccade onset. To do so, the latency between grating onset and saccade onset was recorded on every trial and averaged over the previous 100 trials.

Each experimental session contained a total of 2,400 trials: 1,120 fixation trials, 1,120 saccade trials and 160 control trials, randomly interleaved across 5 blocks. There was a total of 480 trials per block, with a mini-break every 100 trials. Trials were split evenly between fixation and saccade trials (224 trials each) with the remaining allocated to control trials (32 trials).

### 2.3 EEG and eye-tracking pre-processing

EEG and EOG data were recorded at 2048 Hz using a BioSemi system, with 64 active electrodes and 6 ocular electrodes. The continuous EEG and eye-tracking data was pre-processed off-line using MATLAB Version R2020a and EEGLAB toolbox (v2021.0) (Delorme & Makeig, 2004). The data was first down-sampled to 256 Hz and then re-referenced to the mastoids. Eyetracking data were recorded using an Eyelink 1000 eyetracker (SR Research) at 1000 Hz. The eye tracker was calibrated at the start of the experiment and at the beginning of every block. In addition, drift correction was applied at each mini break within a block. The eye-tracking data was synchronised with the EEG using the EYE-EEG toolbox version 0.81 (Dimigen et al., 2011). Saccades were detected using a velocity-based algorithm (Engbert & Kliegl, 2003) within the EYE-EEG toolbox. Successive eye positions were considered saccades if the velocity of the left eye exceeded a threshold of 5 standard deviations (median-based) of all recorded eye velocities (excluding blink intervals) for at least 15 ms. A low threshold was used to ensure the detection of microsaccades which are important for the removal of eye-movement related artifacts in later steps (Dimigen et al., 2009). If the time between two saccadic events was less than 50 ms, only the first saccade was kept, in order to avoid including post-saccadic oscillations as separate saccades (Dimigen, 2020). Following this, saccades were considered as valid if they exceeded an amplitude of 15 dva.

The EEG data was then notch filtered at ∼50Hz to remove electrical artifacts and bandpass filtered between 0.1 and 80 Hz. Automatic data rejection was employed to remove any major artifacts using the Artifact Subspace Reconstruction (ASR) method in EEGLAB. The ASR rejection threshold parameter *k* was set to 15. Bad channels, noted during data collection and confirmed later offline, were spherically interpolated.

To correct for eye movement artifacts in the EEG, we applied independent component analysis (ICA; (Makeig et al., 1996)). To identify eye movement related components, the variance ratio of the component activation during periods of eye movements (blinks and saccades) was compared to that during fixation periods (Plöchl et al., 2012). ICA was performed in a separate pre-processing pipeline containing an additional high-pass filter (Hamming windowed sinc FIR, edge of the passband: 2Hz). It was run on clean continuous data with major movement artifacts removed. The ICA weights were then appended to the corresponding datasets in the original pre-processing timeline and IC activations were recomputed. Components were rejected if the mean variance of activity selected around a saccade (-.02 -.01 ms) was 10% greater than the mean variance during fixation periods (Plöchl et al., 2012; Dimigen, 2020).

For all trial types, if gaze deviated more than 2.5 dva from the current fixation point while the stimulus was on the screen, the trial was removed. Similarly, if the saccade landing was more than 2.5 dva away from the saccade target, the trial was discarded. In addition, only trials in which the physical stimulus was removed from the screen by the time the eyes arrived at the saccade target, were included. Previous studies of transsaccadic fusion (Wolf et al., 1980; Jonides et al., 1982) have been challenged because the results could be explained by lingering visual monitor persistence (Irwin et al., 1983, 1990). This is not the case in the present study as the grey-to-grey time of the monitor used was 1 ms. Additionally, it has been reported that LCD monitors are optimal when visual persistence is a concern given the short rise times (1-6 ms) (Lagroix et al., 2012).

EEG data were epoched for fixation, control and saccade trials separately. Across all trial types, epochs were time-locked to the presentation of the grating. Epochs were extracted from 200 ms before stimulus onset to 500 ms after and were baseline corrected to the mean of the 100 ms period before stimulus onset. Fixation trials were subsequently included in the training set for the main analysis.

### 2.4 Multivariate Pattern Analysis

Epochs from the training set of fixation trials were used to train time-resolved pairwise linear discriminant analysis (LDA) classifiers (Grootswagers et al., 2017) to dissociate the neural activation patterns associated with the presentation of the stimulus at a high and low SF. This was done for each location separately using the amplitude from the 64 electrode channels as input. Each location-specific LDA classifier was tested on independent fixation trials from the trained location and cross-tested on trials from other locations. This was done separately for left and right fixation trials, across all timepoints in the epoch and for each individual participant. Above-chance classification performance indicates that the EEG signal contained information that allowed the classifier to discriminate between a high and low frequency stimulus. The classifiers trained on fixation trials were used to decode SF on control trials and saccade trials.

To examine the time course of SF information availability on saccade trials, the classifiers that were trained to dissociate stimulus SF on fixation trials were tested using data from the peri-saccadic period of saccade trials. This was carried out time-locked to stimulus onset (0 ms to 500 ms after stimulus onset). Only trials in which the saccade occurred in a range between 100-400 ms after stimulus onset were included in the analysis. To understand if features are precisely remapped to the post-saccadic retinotopic position we split saccade trials into ‘congruent’ or ‘incongruent’ depending on the training set location. Congruent trials were those saccade trials in which a stimulus was presented at the same retinotopic position as the position used in the training set, relative to the other fixation point (i.e. 1-5, 2-6, 3-7, 4-8 in Fig. 1a. *right*). This is because the corresponding remapped location overlaps with the trained location. Incongruent trials included the non-remapped location at the same eccentricity. For example, if the classifier was trained on location 1, congruent saccade trials would include location 5 stimuli and incongruent trials would include location 6 stimuli (Fig. 1a. *right*). Classifiers were tested on the corresponding time points used for training i.e., the time diagonal.

### 2.5 Statistical inference

We used Bayes factors (BFs) to determine above-chance decoding and at chance decoding (i.e., null hypothesis) at every timepoint within each of the ten participants using the Bayes Factor R package (Morey & Rouder, 2018) implemented in Python (Teichmann et al., 2021). We set the prior for the null hypothesis at 0.5 (chance decoding) for assessing the decoding results and at 0 for assessing differences across decoding results. A half-Cauchy prior was used for the alternative hypothesis with a medium width of r = √22 = 0.707. Based on Teichmann et al., (2021), we set the standardized effect sizes expected to occur under the alternative hypothesis in a range between -∞ and ∞ to capture above and below chance decoding with a medium effect size (Morey & Rouder, 2018).

BFs larger than 1 indicate that there is more evidence for the alternative hypothesis than the null hypothesis (Dienes, 2011) with a BF greater than 3 considered as “substantial” evidence for the alternative hypothesis i.e., decoding above-chance level. A BF less than 1/3 is considered substantial evidence for the null hypothesis i.e., chance-level decoding (Jeffreys, 1939; 1961).

## 3 Results

### 3.1 Decoding SF across the visual field during stable fixation

Firstly, to test whether we could extract SF information from the EEG signal under stable fixation conditions (i.e. fixation and control trials), we trained classifiers on a subset of EEG fixation trial epochs in which stimuli were presented at either a high or low SF and subsequently tested them on left-out data from the same location. This was done separately for each of the four parafoveal locations (Same condition). Fixation could be on the left or the right of the screen meaning a total of eight classifiers were trained and tested. Classification results were then averaged across all locations.

Subsequently, we tested whether classifiers trained to dissociate SF at one location could perform above-chance when tested on stimuli at varying eccentricities from current fixation. To do this, we ran a cross-location decoding analysis. In this analysis, data from one of the four parafoveal locations were used to train a classifier that was subsequently tested on data from each of the remaining seven locations (Fig. 2a). Classification results were grouped based on stimulus eccentricity from current fixation and averaged within each eccentricity group: Near (∼5 dva), Mid (∼10 dva) and Far (∼20 dva) (Fig. 2b). This allows us to see the degree to which the representation of SF spreads across the visual field. In other words: how well does the representation of SF at one retinotopic position generalise to another?

**Figure 2.**
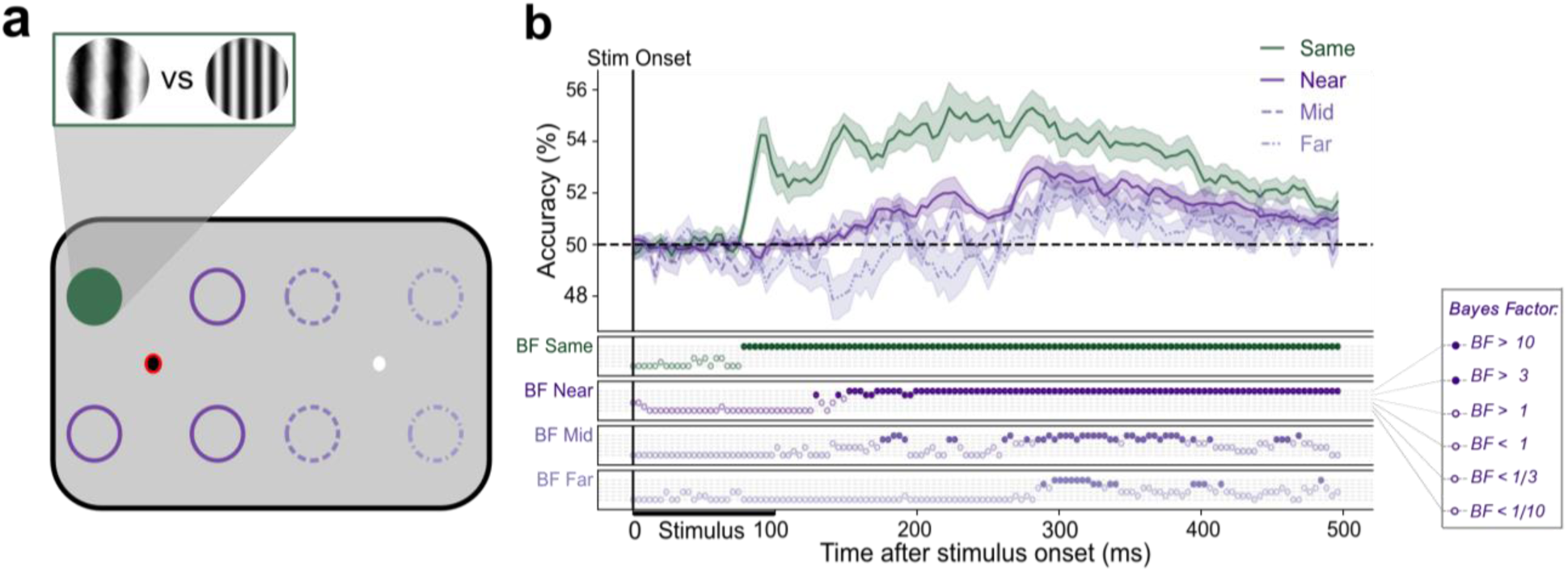
A spatially invariant representation of spatial frequency. Classifiers were trained and tested to distinguish patterns of neural activity evoked by the presentation of a stimulus at different locations on the screen. In panel **a.** the circles represent possible stimulus presentation locations and one example configuration of train and test locations for classification. The dark green circle represents the location that the classifier was trained on, the dark purple circles indicate the near test locations, the medium purple (dashed) indicate the mid test locations and the lightest purple (dot dashed) indicate the far test locations. The red ring signifies current fixation. The gratings illustrate stimuli at low and high SF. **b.** The mean classification over time for fixation trials trained and tested at the same location (green line) and trained on one location and tested on near locations (dark purple line), mid locations (medium purple - dashed) and far locations (light purple – dot dashed). The plotted results reflect the average performance at corresponding training and testing timepoints. All decoding results are averaged over locations at the same eccentricity, across left and right fixation, and subsequently, across subjects. Shaded areas depict standard error of the mean across subjects. The Bayes factors (BF) below the plots indicate the timepoints at which there was substantial evidence in favour of the alternative hypothesis i.e., decoding above-chance (filled circles). Timepoints at which there was not enough evidence or substantial evidence for the null hypothesis, i.e., chance level, are indicated by open circles.

#### 3.1.1 Same location decoding

Using a 5-fold cross-validation procedure, we found that classifiers effectively labelled the test set trials as their correct SF across all locations (Fig. 2b; ‘Same’). Above-chance decoding of SF emerged by 78 ms after stimulus onset and was sustained throughout the entire trial period. This demonstrates that when training and testing at the same parafoveal location, our classifiers could robustly extract SF information from the EEG signal.

#### 3.1.2 Cross-decoding from near eccentric positions

There was substantial evidence for above-chance cross-location decoding of SF at near locations (Fig 2a. *dark purple*). Above-chance decoding emerged by 152 ms, later than when training and testing on the same location (Fig 2b; ‘Near’). This demonstrates that ∼74 ms after SF information is available at the stimulus presentation location, it spreads to adjacent locations at the same eccentricity from fixation. These results indicate that the representation of SF generalises to nearby parafoveal locations after ∼150 ms.

#### 3.1.3 Cross-decoding from mid eccentric positions

We next wanted to investigate whether SF generalises to the rest of the visual field or whether it is constrained to locations near the trained location. The same four parafoveal fixation classifiers were tested on trials in which the stimulus was presented 10 dva away from fixation, in the periphery (Fig. 2a; *medium purple - dashed*). Dissociation between high and low SF first rose above chance at 184 ms post-stimulus onset. (Fig. 2b; ‘Mid’), indicating an emerging generalisation of the SF representation.

#### 3.1.4 Cross-decoding from far eccentric positions

Above-chance cross-decoding of SF at far peripheral locations (20 dva) (Fig 2a. *light purple - dot dashed*) emerged later at 297 ms post-stimulus onset (Fig 2b; ‘Far’). These results indicate an emerging spatially invariant representation of SF that spreads over time from the presented stimulus position across the entire visual field.

### 3.2 Decoding SF at the remapped location with and without a saccade

To examine whether SF is transferred to the remapped position, we ran another cross-location decoding analysis: training on fixation trials and testing on saccade trials. Specifically, we wanted to test if SF at a peripheral location was remapped to its corresponding post-saccadic retinotopic position in the presence of a saccade. Again, four classifiers were trained on SF from parafoveal locations. Importantly, individual classifiers were tested only on trials from the congruent peripheral location. For example, if the classifier was trained on trials from the location to the upper left of current fixation (Fig. 3a, Top panel - filled green circle), it was tested only on trials from the location to the upper left of the saccade target (Fig. 3a, Top panel - orange circle) as these correspond to the pre-saccadic and post-saccadic retinotopic stimulus locations, respectively (see section 2.4 for more details). The same cross-location decoding was done for control trials with no saccade (Fig 3a. Bottom panel - purple circle).

**Figure 3.**
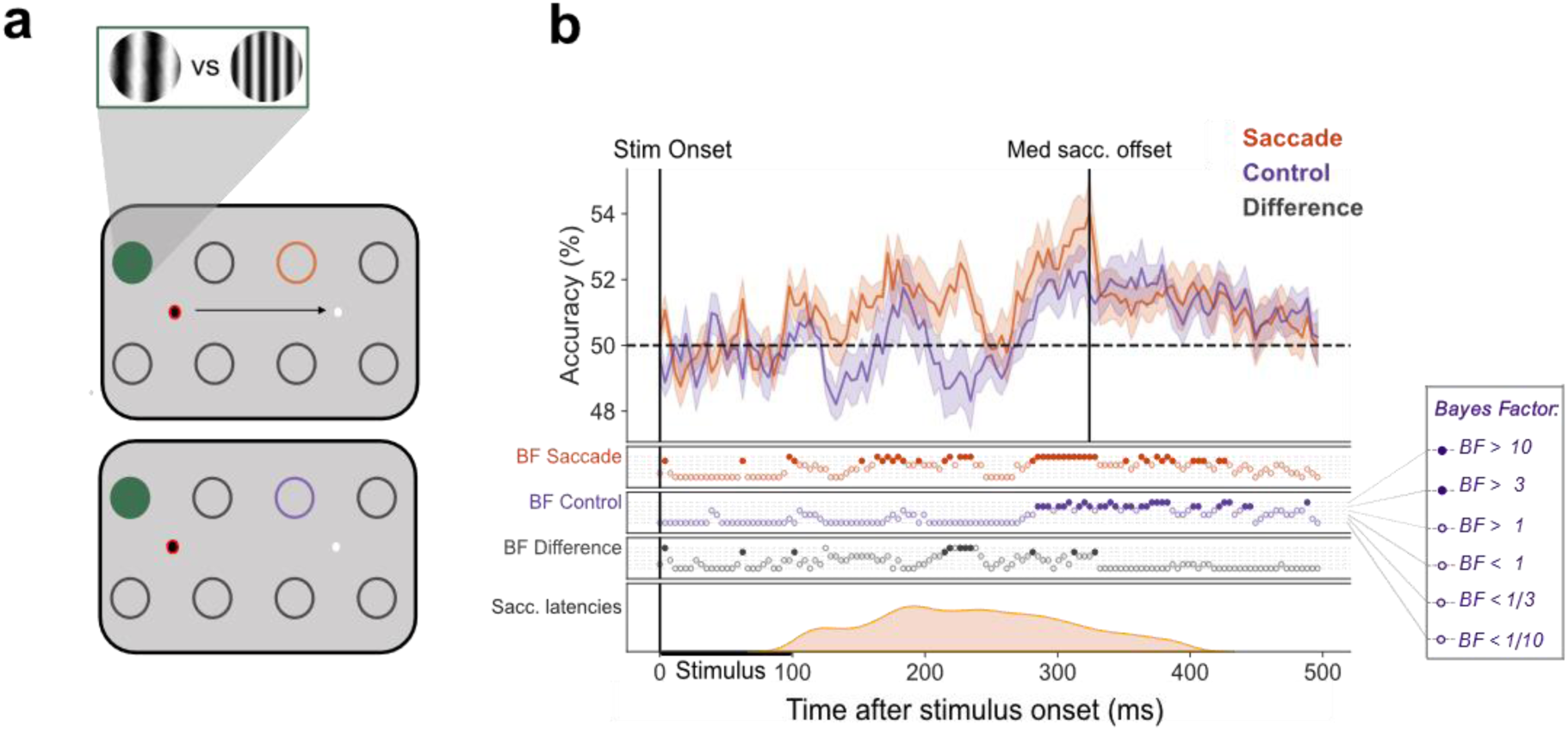
Earlier decoding of SF at the remapped location with a saccade. This figure illustrates the difference between decoding at the remapped location with and without a saccade. In panel **a** the circles represent possible stimulus presentation locations and an example configuration of train and test locations for classification of saccade trials (top panel) and control trials (bottom panel). The green filled circle represents the location on which the classifier was trained and the orange and purple circles signify the test locations for saccade and control trials, respectively. The red ring signifies current fixation. The gratings illustrate stimuli at low and high SF**. b.** The mean classification over time for classifiers trained on fixation trials and tested on either saccade trials (orange) or control (purple) trials. The plotted results reflect the average performance at corresponding training and testing timepoints. All decoding results are averaged over classifiers, across left and right fixation, and subsequently, across subjects. Shaded areas depict standard error of the mean across subjects. The Bayes factors (BF) below the plots indicate the timepoints at which there was substantial evidence in favour of the alternative hypothesis i.e., decoding above-chance (filled circles). Timepoints at which there was not enough evidence or substantial evidence for the null hypothesis, i.e., chance level, are indicated by open circles. The distribution of saccade onset times is plotted below.

As shown in the previous analyses: under stable fixation SF decoding generalises across the entire visual field. We therefore expect to find SF information at the ‘remapped’ position even in the absence of a saccade. Notably, however, the temporal availability of SF information at the remapped location should gain a predictive nature under active vision. In other words, the latency of above-chance decoding of SF at the remapped location should be earlier for saccade trials compared to control trials in which no saccade is performed (but stimulus presentation is otherwise equivalent).

From 164 ms post-stimulus onset in saccade trials, the classifier was able to predict above-chance which SF was presented in the periphery. This is in contrast to decoding of peripheral trials in the absence of a saccade (i.e., control) where peripheral SF could be reliably decoded only 100 ms later, at 285 ms post-stimulus onset (Fig 3b). Substantial differences in the decoding time courses between conditions emerged at 219 ms post-stimulus onset and was maintained until 238 ms. These results indicate that in the presence of a saccade, information about SF is available at the remapped location earlier than during stable fixation. This finding is consistent with predictive remapping of SF as a result of the saccade.

### 3.3 Remapping of SF is spatially coarse

The previous analysis showed initial evidence of predictive remapping of peripheral stimulus SF to the remapped location. Next, we wanted to determine the extent to which this predictive representation is spatially precise. To determine whether SF information is transferred uniquely to the remapped location in the presence of a saccade, we used the same trained fixation classifiers to test peripheral stimuli presented at congruent (Fig 4a.; light orange - solid) and incongruent locations (Fig 4a; dark orange - dashed).

**Figure 4.**
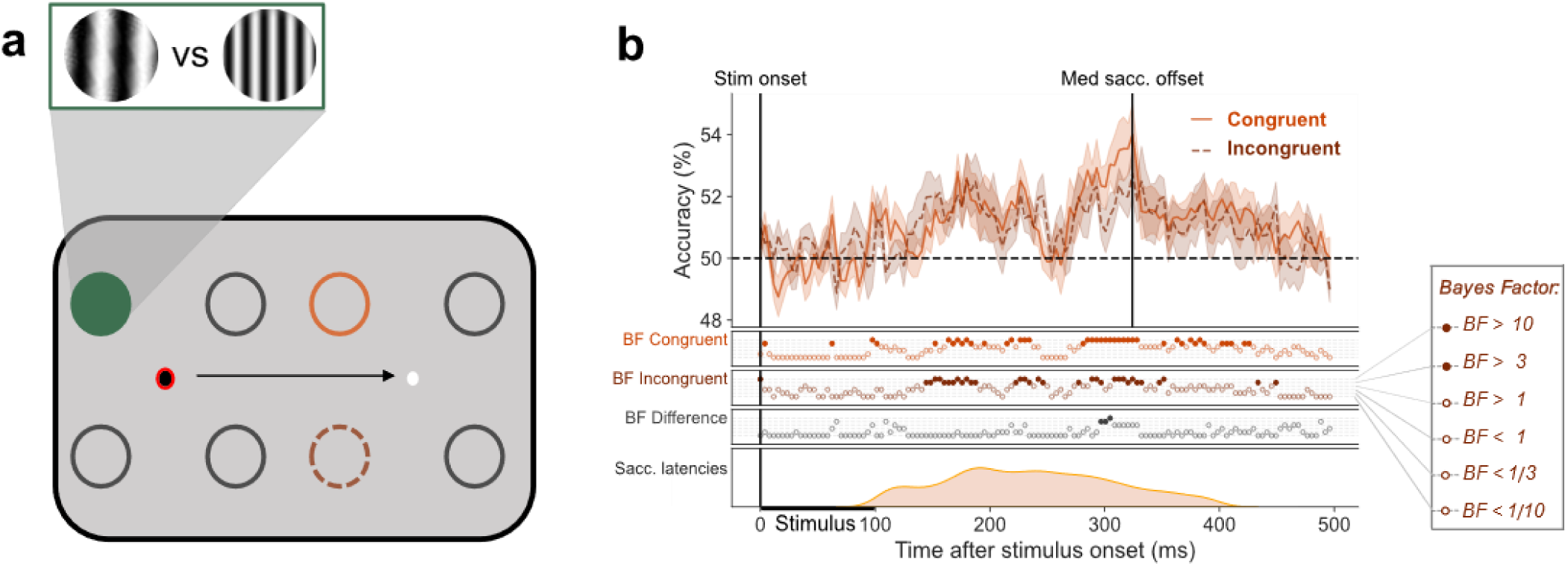
No spatially precise remapping of SF. **a.** Classifiers were trained on high vs low frequency stimuli presented at each of the four locations around current fixation (red ring) producing four training sets. Illustrated is an example configuration of train and test locations for classification. The green filled circle represents the location that the classifier was trained on. Each training set was paired with two test sets. One test set in which stimuli were presented at a congruent location (light orange – solid line) and another in which stimuli were presented at an incongruent location (dark orange – dashed line) in the periphery. The black arrow represents the saccade. The gratings illustrate stimuli at low and high SF**. b.** The mean classification over time for congruent trials (light orange – solid line) and incongruent trials (dark orange – dashed line) in saccade trials. The plotted results reflect the average performance at corresponding training and testing timepoints. Shaded areas depict standard error of the mean across subjects. All decoding results are averaged across left and right fixation, and subsequently, across subjects. The Bayes factors (BF) below the plots indicate the timepoints at which there was substantial evidence in favour of the alternative hypothesis i.e., decoding above-chance (filled circles). Timepoints at which there was not enough evidence or substantial evidence for the null hypothesis, i.e., chance level, are indicated by open circles. The distribution of saccade onset times is plotted below.

SF could be decoded from both the congruent and incongruent locations with no reliable evidence for any difference between decoding at these locations (Fig. 4b). This suggests that the ability to decode SF from peripheral positions cannot be explained by a precise transfer of stimulus features to the post-saccadic retinotopic position. Rather, decoding is more spatially coarse such that each parafoveal classifier (e.g., Fig 4a; green – filled circle) can decode SF from all peripheral positions. Thus, we can conclude from this analysis that while the saccade allows predictive anticipation of peripheral stimulus features, this response does not contain information about the post-saccadic retinotopic position of the stimulus.

### 3.4 Representation of peripheral spatial frequency at parafoveal positions

After establishing the absence of precise spatial remapping, we wanted to characterise the representation of SF during a saccade more generally. To do this we trained classifiers on SF at the four parafoveal locations using fixation trial data, (Fig 5a; Example training location shown by filled green circle) and cross-tested them on Near (Fig 5a; dark purple), Mid (Fig 5a; light orange – solid line) and Far trials (Fig 5a; dark orange – dashed line) during the saccadic period. The classification scores of the classifiers were averaged within each eccentricity group. This gives us an understanding of when SF at different eccentricities becomes available at positions around fixation during the peri-saccadic period.

**Figure 5.**
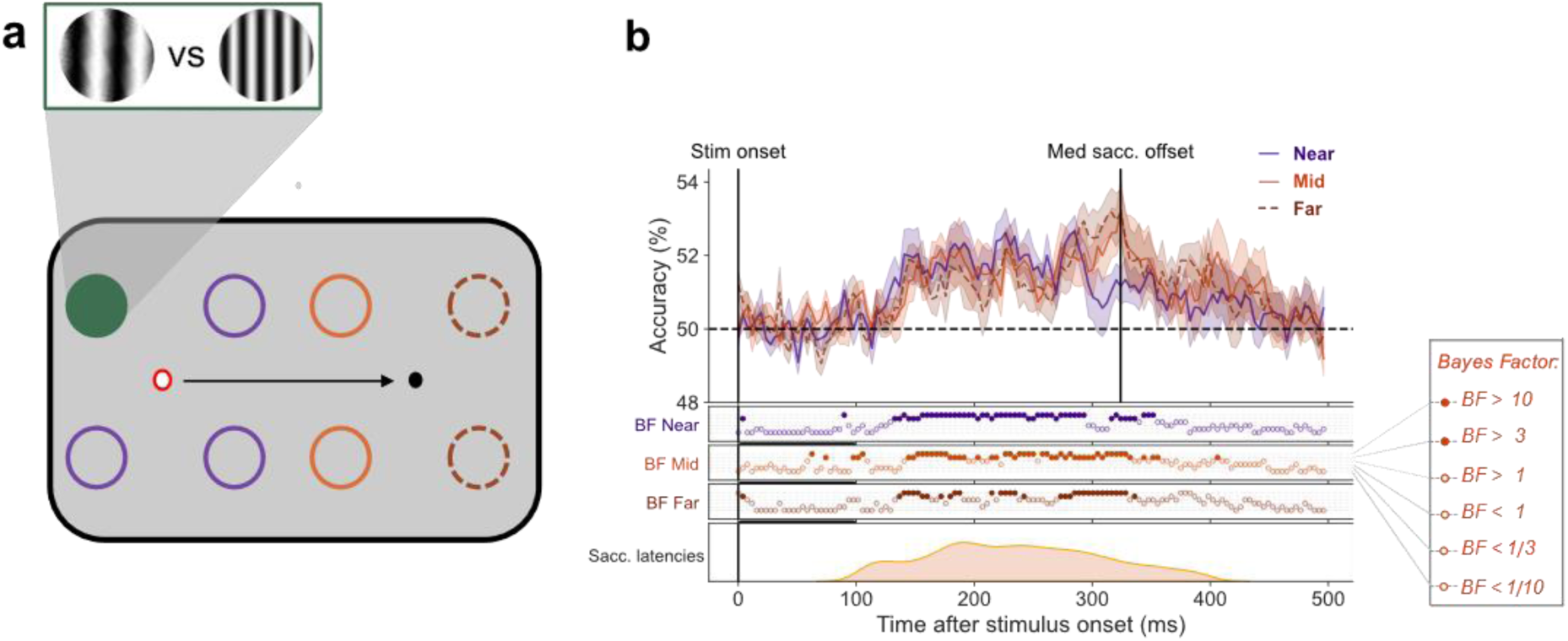
Similar decoding time course across eccentricities in the presence of a saccade. **a.** Classifiers were trained on high vs low frequency stimuli presented at each of the four locations around current fixation (red ring) producing four training sets. Illustrated is an example configuration of train and test locations for classification. The filled green circle represents the location that the classifier was trained on, the dark purple (solid line), light orange (solid line) and dark orange (dashed line) indicate the near, mid and far test locations (during saccade trials), respectively. The red ring signifies current fixation. The gratings illustrate example stimuli at low and high SF. The black arrow represents the saccade. **b.** The mean classification over time for classifiers trained on fixation trials and tested on Near (dark purple), Mid (light orange – solid line) and Far locations (dark orange – dashed line). The plotted results reflect the average performance at corresponding training and testing timepoints. All decoding results are averaged over eccentricity, across left and right fixation, and subsequently, across subjects. The Bayes factors (BF) below the plots indicate the timepoints at which there was substantial evidence in favour of the alternative hypothesis i.e., decoding above-chance (filled circles). Timepoints at which there was not enough evidence or substantial evidence for the null hypothesis, i.e., chance level, are indicated by open circles. The distribution of saccade onset times is plotted below.

Above-chance decoding of SF for Near, Mid and Far stimuli first rises above chance at 132, 145 and 137 ms, respectively (Fig 5b), with no substantial differences between the decoding time courses. This indicates that during the peri-saccadic period at ∼140 ms post stimulus onset, across all stimulus eccentricities, the representation of SF is similar to the parafoveal representation of SF during stable fixation. This is a considerably shorter decoding latency than that found for peripheral stimuli under stable fixation at both Mid (184 ms) and Far (287 ms) positions and slightly shorter for Near positions (152 ms). Interestingly, the saccade seems to remove latency differences in the spread of SF across eccentricities, giving access to SF information at a similar rate.

## 4 Discussion

In this study, we used multivariate pattern analysis (MVPA) of EEG data to understand if stimulus features are predictively remapped. This method provides a spatiotemporally precise picture of the neural representation of SF, allowing us to characterise the spread of SF information across the visual field during stable fixation. In turn, we determined the extent to which saccade planning provided predictive access to peripheral SF by comparing decoding latencies across fixation and saccade conditions.

We first show an emerging position invariant representation of SF during stable fixation. We found that, despite training a classifier on SF at parafoveal locations, SF could be decoded from positions up to 20 dva in the periphery. The timing of above-chance decoding was later as eccentricity increased, providing a time course for the spreading of SF information across the visual field. Next, we found that the decoding latency of peripheral SF was accelerated in the peri-saccadic period compared to stable fixation, initially suggesting predictive remapping of SF. Interestingly however, peripheral stimulus SF could be decoded from all parafoveal positions, not just the remapped location, at a similar latency. This indicates that the predictive representation of SF is unbound to a specific retinotopic position. Overall, these results support a predictive representation of peripheral SF due to a saccade which precedes the emergence of a position invariant representation.

### The spread of spatial frequency across the visual field

We found that the representation of SF spreads gradually over the visual field. This is consistent with stimulus features that are initially processed only by neurons with receptive fields responsive to the stimulus location, but later in a more global fashion across the visual cortex. The early above-chance peak (78 ms), unique to same location decoding likely signifies the initial retinotopic coding of the stimulus in which SF is bound to its retinotopic position. Once visual input has moved through low-level and mid-level extrastriate areas (V2-V5), stimulus activity begins to spread beyond strictly retinotopic areas to higher level areas with larger receptive fields (Wurtz & Kandel, 2000).The ability to decode SF at spatial locations adjacent to the trained location (Near eccentricity), starting at ∼150 ms after stimulus onset, is consistent with the visual information reaching higher visual areas with larger receptive fields, such as the inferior temporal cortex (Richmond & Wurtz, 1983;Keysers et al., 2001 Hung et al., 2005) that are invariant to changes in object position (Logothetis et al., 1995). The timing of this spatially invariant response also aligns well with human EEG (Liu et al., 2009) and MEG (Fabius et al., 2020) recordings Above-chance decoding of SF at peripheral eccentricities emerged at 184 ms (Mid) and 297 ms (Far) post-stimulus onset providing further evidence for the spatially global spread of SF. These results demonstrate that despite large retinotopic eccentricity differences, after ∼290 ms the stimulus-driven EEG activity is similar across all eccentricities. This could be explained by feature based attention (FBA), which modulates the firing rate of a neuron tuned to a particular stimulus feature despite it not being directly driven by a stimulus in its RF (Serences & Boynton, 2007). Previous time-resolved research examining the spread of FBA has only used a single stimulus eccentricity, usually positioned within ∼7 dva of fixation with FBA effects emerging at ∼200-280 ms post stimulus onset (Anllo-Vento & Hillyard, 1996; Stoppel et al., 2012; Bartsch et al., 2015, 2017, 2018). We extend this by combining EEG, which provides high temporal resolution, with MVPA, allowing us to pinpoint when neural representations overlap in time, ultimately providing a temporally resolved delineation of the spread of FBA.

### Saccades allow predictive access to peripheral spatial frequency

In line with previous reports of feature remapping, we found that SF could be decoded at the remapped position during the peri-saccadic period. Importantly, SF information at that location was available earlier on saccade trials (∼160 ms) compared to the corresponding location on control trials (∼285 ms). This can be considered a predictive response as it occurs at a shorter latency than the typical visual response (Merriam et al., 2007) (i.e., cross-decoding the same location in the absence of a saccade). If the availability of SF at the remapped position was the manifestation of saccade-independent feedback processes, such as FBA, we would expect above-chance decoding to emerge at a similar time to control trial decoding given that stimuli overlapped in their retinotopic position. This was not the case.

Critically however, this predictive response was not specific to the remapped location. Instead, the upcoming saccade seems to facilitate predictive access to SF information at all stimulus positions. Unlike under stable fixation, where decoding latency parametrically increased with eccentricity, in the presence of a saccade stimulus SF could be decoded across the visual field at the same time (∼140 ms) and importantly, before the earliest cross-decoding during fixation (152 ms).

The spatially coarse remapping of SF is difficult to reconcile with the precise nature of forward saccadic remapping (Collins et al., 2009), however may be explained by convergent remapping, where receptive fields converge towards the saccade target (Tolias et al., 2001; Zirnsak & Moore, 2014). Zirnsak and Moore (2014) reported that when decoding from FEF neurons around the time of a saccade, stimuli were miscategorised with an average error of 7 dva. This aligns with our finding that in saccade trials, SF at both Mid and Far positions followed a similar decoding time course, suggesting overlapping neural representations. Both Mid and Far stimuli were positioned at the same distance from the saccade target (5.05 dva). The close proximity of our stimuli to the saccade target and the predictive shift of population RFs (pRF) towards the saccade endpoint, as stated in convergent remapping theories, may mean that mid and far stimuli were similarly encoded. Stimuli at both Mid and Far positions may have fallen within the shifted pRF, evoked a similar neural response despite different retinotopic coordinates. How exactly convergent remapping contributes to stable vision or the continuity of features across saccades is not known (Marino & Mazer, 2016).

### Could foveal feedback explain predictive representation of spatial frequency?

Classifiers were trained on SF at four possible retinotopic positions (parafoveal locations) however, all training stimuli extended into the foveal region, likely eliciting somewhat overlapping representations. Recent work investigating foveal feedback in the pre-saccadic period found that during saccade preparation, a transient connection is opened between the current and future foveal positions. Using a behavioural detection task, Kroell and Rolfs (2022) found that participants had a higher hit rate for foveal probes when they matched the orientation of the saccade target. Importantly, this effect extended to stimuli up to 6.4 dva around fixation. Interestingly, participants had more congruent false alarms meaning that the saccade target feature was boosted at foveal locations causing participants to ‘see’ it in the noise even when no probe was presented. This suggests an enhancement of the relevant features at foveal locations.

Similarly, we found that peripheral SF could be decoded from the four locations around fixation in anticipation of a saccade. While in our study the stimuli were not the target of the saccade, stimulus SF information may still have been fed back to foveal retinotopic cortex. Previous research, outside the saccade literature, demonstrates that relevant peripheral information is represented at foveal retinotopic cortex (Williams et al., 2008; Fan et al., 2016). A shown by earlier decoding at locations extending into the fovea when a saccade was imminent, the saccade seems to accelerate this feedback process. This supports the idea that feedback connections to foveal retinotopic cortex function to predict stimulus features during saccade preparation (Kroell & Rolfs, 2022).

To better discern the mechanism underlying the predictive neural response reported here, future studies should ensure that training locations are well separated such that stimuli fall either within the parafoveal or foveal region. The parafoveal positions should correspond to the remapped positions of the peripheral stimuli. By more precisely separating training locations, one could check whether peripheral stimulus features are predictively represented only at the exact remapped position, only at the fovea or more generally across all trained locations.

### Saccade onset time does not modulate latency of decoding

Despite predictive decoding of SF due to the presence of a saccade, the latency at which SF information spread to parafoveal positions was not modulated by the timing of saccade onset. We split saccade trials into different bins depending on when the saccade occurred after stimulus onset (100-400 ms; Fig. 6) and found that above-chance decoding of peripheral SF at parafoveal positions emerges around ∼150 ms, similar to the latency found when

**Figure 6.**
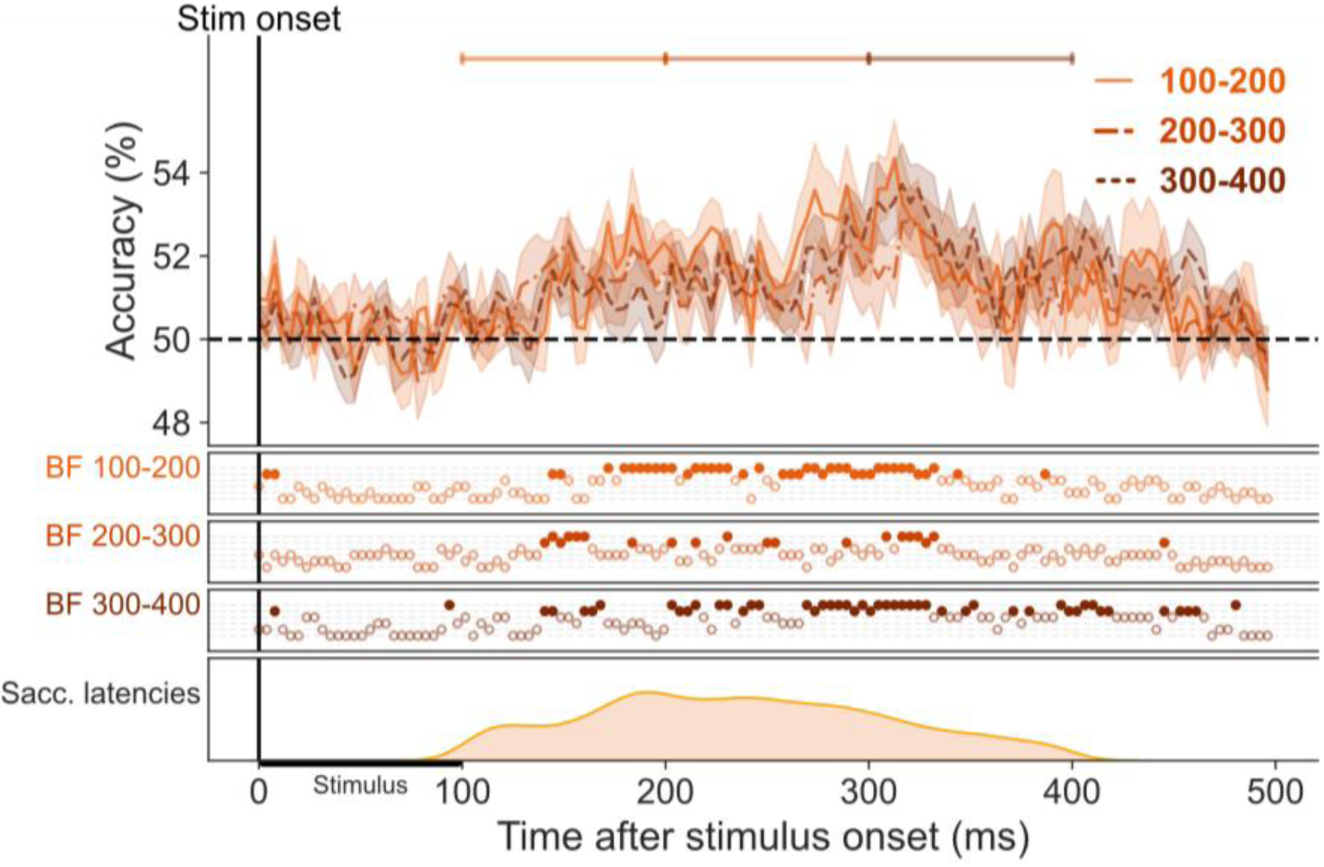
No substantial difference in decoding time course across saccade latencies. The mean classification over time for classifiers trained on fixation trials and tested on saccade trials, split into different bins depending on the latency between stimulus onset and saccade onset. The plotted results reflect the average performance at corresponding training and testing timepoints. All decoding results are averaged over locations, across left and right fixation, and subsequently, across subjects. The Bayes factors (BF) below the plots indicate the timepoints at which there was substantial evidence in favour of the alternative hypothesis i.e. decoding above-chance (filled circles). Timepoints at which there was not enough evidence or substantial evidence for the null hypothesis, i.e. chance level, are indicated by open circles.

combining saccade trials across all bins (main results). This is strange, as, if saccade preparation is driving predictive responding, one would expect saccade timing to dictate when this information becomes available. The fact that we see similar results across all bins even those outside the normal saccade preparation range may be explained by the sudden onset of our target stimulus. Increased saccade latencies have been reported when a peripheral stimulus needs to be ignored in order to accurately perform the saccade. While our data cannot reveal if stimulus onset affected saccade latencies as we lack an informative control, previous research has shown that even if the saccade has been prepared, the onset of a peripheral stimulus can prolong the latency of its onset (Dalmaso et al., 2020). This suggests that in the current setup, saccade onset time is not a reliable marker of when saccade preparation begins. Saccade preparation and its associated predictive processes may have begun well before saccade onset but actual initiation of the saccade is delayed. This may explain the predictive decoding of SF across all bins. Still, the seeming independence of SF decoding from saccade timing is in stark contrast to spatial information around the time of saccades. Under the same experimental conditions, we found that the remapped location could only be decoded when the stimulus appeared immediately before saccade onset (Moran et al. 2024). Further research is needed to understand these timing differences. Additionally, it is important to understand how spatial remapping and the spatially coarse remapping of stimulus features reported here work in tandem to produce visual stability.

### Feature based attention on an unattended feature

The pervasive representation of SF across the visual field in the current experiment is somewhat surprising given that it was task irrelevant. We chose a behavioural task that was orthogonal to the studied feature, to assess automatic remapping of stimulus features and remove any confound of task relevance. However, there is substantial evidence from both single-cell animal recordings (Katzner et al., 2009) and human neuroimaging studies (O’Craven et al., 1999; Schoenfeld et al., 2003; Ernst et al., 2013a) to suggest that attention to a single feature can spread to a secondary feature if the features are part of the same object.

For example, blood oxygenation level-dependent (BOLD) responses related to stimulus motion were enhanced even when participants were cued to attend to the colour of moving dots (Ernst et al., 2013a). Not only does attention spread to the secondary feature, but the task-irrelevant feature can be enhanced across the entire visual field, similar to results reported for attended features (Scholl, 2001). These findings, as well as our own results, support object-based theories of attention in which all features of an attended object are automatically selected for processing, regardless of their task-relevance (Duncan, 1984; Scholl, 2001).

### Consequences of spatially invariant representation of spatial frequency

Our results indicate that a global representation of SF emerges after ∼250 ms, potentially indicating that precise remapping of features to the post-saccadic retinotopic location is less urgent than remapping of spatial location. The pervasive representation of SF across the entire visual field calls into question the validity of previous reports of feature remapping. For example, studies that have examined TAE under saccade conditions (Melcher, 2007 & He et al. 2018), found that an adaptor, when presented at the post-saccadic retinotopic location of a test stimulus, induced a TAE effect. If, as demonstrated here, features are processed to some degree in a position invariant manner, then the results reported previously may not be due to a spatially precise remapping of features. This competing hypothesis is particularly likely in studies in which participants have a long stimulus exposure time, giving the spatially invariant representation time to develop. In such a case, neurons tuned to a specific feature are activated across the visual cortex with the adaptor stimulus and so the TAE should be found regardless of its retinotopic position. This is in line with a study that investigated the spread of a motion after effect (MAE) across the visual field. They found that an adaptor presented at one location induced an MAE evenly across all other locations tested (Liu & Mance, 2011). Similar results were found when looking at the TAE (Liu & Hou, 2011). This was in the absence of any saccade and so speaks to the global nature of feature-based attention.

## Conclusion

We demonstrate an emerging position invariant representation of SF. In addition, we show spatially coarse predictive remapping of SF in the presence of a saccade, whereby access to peripheral stimulus feature information is accelerated. Importantly, this predictive representation is not isolated to the remapped location. This predictive process that allows the visual system earlier access to peripheral stimuli may allow the visual system to maintain a stable representation of the environment and facilitate smooth perception despite frequent eye movements. This is complemented by a spatially invariant representation of stimulus features on saccade landing, creating a robust and flexible system for visual processing.

